# miR-544-3p mediates arthritis pain through regulation of FcγRI

**DOI:** 10.1101/2021.06.13.448256

**Authors:** Yan Liu, Sang-Min Jeon, Michael J. Caterina, Lintao Qu

## Abstract

Chronic or episodic joint pain is a major symptom in rheumatoid arthritis (RA) and its adequate treatment represents an unmet medical need. However, the cellular and molecular mechanisms underlying RA pain remain elusive. Non-coding microRNAs (miRNAs) have been implicated in the pathogenesis of RA as negative regulators of the stability or translation of specific target mRNAs. Yet, their significance in RA pain is still not well defined. We and other groups recently identified neuronally expressed FcγRI as a key driver of arthritis pain in mouse RA models. Thus, we tested the hypothesis that miRNAs that target and regulate neuronal FcγRI attenuate RA pain. Here, we show that miR-544-3p was robustly downregulated whereas FcγRI was significantly upregulated in the dorsal root ganglion (DRG) in mouse RA models. Intrathecal injection of miR-544-3p mimic attenuated established mechanical and heat hyperalgesia in a mouse model of collagen II-induced arthritis (CIA). Moreover, this effect was likely mediated, at least in part, by FcγRI since miR-544-3p mimic downregulated FcγRI in the DRG during arthritis and genetic deletion of FcγRI produced similar antihyperalgesic effects in the CIA model. This notion was further supported by a dual luciferase assay showing that miR-544-3p targeted FcγRI by directly binding to its 3’UTR. In addition, FcγRI expression in DRG neurons in vitro was downregulated by miR-544-3p mimic and upregulated by miR-544-3p inhibitor. In naïve mice, miR-544-3p mimic alleviated acute joint pain hypersensitivity induced by IgG immune complex (IgG-IC), whereas miR-544-3p inhibitor potentiated the pro-nociceptive behavioral effect of IgG-IC. These findings suggest that miR-544-3p causally participates in the maintenance of arthritis pain by targeting neuronal FcγRI, and thus define miR-544-3p as a new potential therapeutic target for treating RA pain.

## Introduction

Rheumatoid arthritis (RA) is a common chronic autoimmune disease with high morbidity and mortality that primarily affects joints ^1^. Joint pain is a cardinal clinical feature of RA that significantly impacts quality of life ^2^. Although RA pain is often thought to be of inflammatory origin, pain outcomes in RA can be poor despite optimal control of inflammation with current biologic therapies ^3^. Therefore, new targeted therapies to effectively manage RA pain are urgently needed.

MicroRNAs (miRNAs) are small noncoding RNAs (∼22 bp) that derive from distinctive hairpin precursors.^4^ As master regulators of gene expression, mature miRNAs degrade mRNA or suppress protein translation by binding target mRNAs at complementary sites in their 3’ untranslated regions (UTR) reminiscent of the RNA induced silencing complex ^5^. Given that both the sequence of miRNAs and their target sites in mRNAs are extensively conserved across mammals ^6^, studies of microRNAs in non-human disease models are of substantial potential clinical relevance ^7^. A variety of miRNAs have been implicated in the pathogenesis of RA ^8^. miR-155 deficient mice showed resistance to collagen II-induced arthritis (CIA) with profound suppression of Th17 cells and antibody responses ^9,10^. miR-106b promoted the development of joint inflammation and bone damage in the CIA model ^11^. In RA patients, serum miR-22 and miR-103a might predict RA development in susceptible individuals (pre-RA) while serum miR-223 levels have been associated with RA activity and disease relapse ^12^. These studies suggest that miRNAs might contribute to arthritis pathogenesis in multiple ways. Yet, none of these studies specifically addressed the potential role of miRNAs in RA pain, per se.

Primary sensory neurons with cell bodies in trigeminal ganglia or dorsal root ganglia (DRG) represent the primary origin of nociception, and are involved in both initiation and maintenance of pathological pain states^13^. Growing evidence suggests that miRNAs expressed in the DRG function as crucial pain modulators via the regulation of pain associated genes^14,15^. miRNA cluster miR-17-92 (including miR-17, miR-18, miR-19a, miR-19b and miR-92) were up-regulated in the DRG in neuropathic pain models and in turn down-regulated potassium channels including Kcna1, Kcna4, Kcnc4, Kcnd3 and Kcnq5^16^. Two α subunits of voltage-gated sodium channels (i.e. Scn3a and Scn9a) were shown to be targeted by miRNAs (miR-30b, miR-96 and miR-183) down-regulated in the DRG following nerve injury^17–20^. In addition, miR-7a was downregulated in the DRG in the rat spinal nerve ligation model and overexpression of miR-7a attenuated neuropathic pain through the downregulation of a β2 subunit of voltage-gated Na^+^ channels^13^. In the Complete Freund’s adjuvant (CFA)-induced chronic inflammatory pain model, miR-23 negatively regulated the expression of µ opioid receptors in DRG neurons^21^. A recent study revealed that miR-143-3p was downregulated in the DRG in the CIA model ^22^. Moreover miR-143-3p can target and regulate the expression of multiple pain associated genes in DRG neurons, including prostaglandin-endoperoxide synthase 2 (Ptgs2), MAS related G protein coupled receptor family member E (Mrgpre) and tumor necrosis factor (Tnf) ^22^.

FcγRI, an immune receptor for IgG-immune complex (IgG-IC), is typically expressed in immune cells and is critical to the regulation of immunity^23,24^. Recently, we and other groups revealed that FcγRI was also expressed in a subset of joint sensory neurons^25,26^. Moreover, upregulated neuronal FcγRI signaling mediated hyperactivity of joint sensory neurons and chronic arthritis pain through a mechanism independent of inflammation^25,26^. Although the functional upregulation of FcγRI in DRG neurons during arthritis may be due in part to augmented FcγRI transcription by proinflammatory mediators, it is also possible that FcγRI expression is upregulated posttranscriptionally via reductions in miRNA-mediated suppression of protein translation or mRNA stability. If so, downregulation of FcγRI expression by specific miRNAs might provide a powerful means of disrupting pain in the setting of arthritis. However, little is known about whether and which miRNAs contribute to the development and maintenance of arthritis pain via the regulation of FcγRI. In this study, we identified miR-544-3p as a crucial miRNA that targets and binds to the 3’UTR of *Fcgr1*. We also showed that miRNA-544-3p downregulation promoted acute joint pain hypersensitivity induced by IgG-IC and chronic arthritis pain by disinhibiting the expression of FcγRI in the DRG.

## Material and methods

### Animals

Adult mice of both genders that were 2 to 4 months old and 20 to 30 g body weight were used for all experiments. Animals were housed under a 14-hour light/10-hour dark cycle with ad libitum access to food and water. Adult DBA1/1J and C57BL/6 were purchased from the Jackson Laboratories (Bar Harbor, ME). Breeders of global *Fcgr1^-/-^* mice were provided by Sjef Verbeek (Leiden University Medical Center, Leiden, The Netherlands). *Fcgr1^-/-^* /DBA/1j mouse littermates were generated by interbreeding of heterozygotes on the DBA/1j background. All animal experimental procedures were approved by the Institutional Animal Care and Use Committee of Johns Hopkins University School of Medicine and were in accordance with the guidelines provided by the NIH and the International Association for the Study of Pain.

### Antigen-induced arthritis (AIA)

AIA was induced in male and female mice on a C57BL/6 background using the antigen methylated bovine serum albumin (mBSA, Sigma, St Louis, MO) as described previously^26^. Briefly, mice were sensitized with 500 µg of mBSA in 200 µl of an emulsion containing 100 µl saline and 100 µl CFA (1 mg/ml; Sigma, St Louis, MO) and delivered by subcutaneous (s.c.) injection to the caudal back skin using a sterile syringe and 25 G needle. Mice were boosted with the same preparations on day 7. Immunized mice were challenged on day 21 by intraarticular (i.a.) injection of mBSA (60 µg; 20 µl in saline) or saline alone (vehicle) to the knee cavity. To induce a flare-up reaction, mice were rechallenged following the same procedure on day 42^27,28^.

### Collagen-induced arthritis (CIA)

CIA was induced in adult mice of both genders on a DBA/1J background using bovine collagen II (Chondrex Inc, Woodinville, WA) ^29,30^. On day 0, adult mice were immunized by intradermal injection of 100 µg of bovine type II collagen that was emulsified in 100 µl of emulsion containing 50 µl acetic acid (0.01 M) and 50 µl CFA (1 mg/ml Myobacterium tuberculosis; Chondrex Inc, Woodinville, WA) at the base of the tail. Twenty-one days after primary immunization, a booster injection with 100 µl of an emulsion of bovine collagen II (100 µg, 50 µl in 0.01 M acetic acid) and Incomplete Freund’s Adjuvant (IFA, 50 µl; Chondrex Inc, Woodinville, WA) was administered intradermally at a different location of the mouse tail. The control group received the same injection but without collagen II. Arthritis score was used as an index of inflammation and disease activity, and assigned based on a scoring protocol in which each swollen or red phalanx was given 0.5 point and 1 point per toe. A red or swollen knuckle was given 1 point, a red or swollen footpad was given 1 point and a swollen ankle and/or wrist was given 5 points. The maximum score for each paw was 15 points, resulting in a maximum possible score of 60 points per mouse^31^.

### Behavioral tests

All behavioral measurements were performed on awake, unrestrained and age-matched littermates (2-3 months) by experimenters blinded to genotype and treatment. Primary mechanical allodynia in ankle joints was measured by applying ascending forces to the ankle with calibrated electronic blunt forceps (Bioseb, Pinellas Park, FL). The cutoff force was set at 350 g to avoid joint damage. The mechanical threshold was defined as the force at which the mouse withdrew its hindlimb forcefully or vocalized ^26^. The mechanical threshold in the joint was averaged over three measurements obtained at intervals of at least 5 min. Secondary mechanical hyperalgesia in the glabrous skin of hind paws was evaluated using von Frey monofilaments of different forces ranging from 0.04 to 1.0 g. Paw withdrawal responses to ten applications of each filament were counted. Secondary thermal hyperalgesia in the glabrous hind paw skin was assessed by measuring withdrawal latency to noxious heat stimuli delivered using a radiant heat source (Plantar Test Apparatus, IITC Life Science Inc., Woodland Hills, CA). The cutoff latency was set at 15 s. Heat response latencies were averaged over three measurements obtained at intervals of at least 3 min.

### Preparation and intra-articular injection of IgG immune complex (IgG-IC)

IgG-IC was formed by incubation of mouse collagen II (1 mg/ml; Chondrex Inc, Woodinville, WA) and anti-mouse collagen II antibody cocktail (1 mg/mL; Chondrex Inc, Woodinville, WA) at a mass ratio of 1:1 for 1 hour at 37°C ^25^. To evaluate the effect of miR-544-3p mimic on IgG-IC induced acute articular hypernociception, 10 μl of IgG-IC at a dose of 100 μg/ml was injected into the right ankle joint cavity of naive mice. For the experiment on miR-544-3p hairpin inhibitor, IgG-IC (10 μg/ml; 10 μl) was chosen for i.a. injection. Pain-related behaviors were measured 1-5 hrs after IgG-IC injection.

### In situ hybridization (ISH) and immunohistochemistry (IHC)

Anesthetized mice were transcardially perfused with diethylpyrocarbonate pre-treated PBS, followed by 4% paraformaldehyde (PFA). L4-L5 DRGs were dissected, cryopreserved in 30% sucrose andcryosectioned at 12 µm. ISH for *Fcgr1* mRNA was performed as described previously^26^. SP6 transcribed antisense and T7 transcribed sense control probes were synthesized from mouse Fcgr1 (NM_010186) cDNA clone (MR225268, OriGene) using 1 set of primers (forward, 5′-ATTTAGGTGACACTATAGAATCCTCAATGCCAAGTGACCC-3′; reverse, 5′-GCGTAATACGACTCACTATAGGGCGCCATCGCTTCTAACTTGC-3′). The probes were then labeled using a digoxigenin (DIG) RNA labeling Kit (Roche Diagnostics Corp., Indianapolis, IN). After pre-hybridization, sections were hybridized with probes (2 ng/µl) at 60°C overnight and washed with 0.2x SSC at 62°C. *Fcgr1* mRNA signal was then detected using sheep anti-DIG antibody (1:500; Roche Diagnostics Corp., Indianapolis, IN) followed by the secondary antibody (Donkey anti-sheep IgG Alexa 488, 1:200; Abcam).

For miR-544-3p, ISH was performed using a miRCURY LNA miRNA ISH Optimization Kit (Qiagen, Germantown, MD) accord to the manufacturer’s instructions. Briefly, target miRNAs were hybridized with 40 nM of specific DIG-labelled miRCURY probes or scramble probes diluted in hybridization buffer (Qiagen, Germantown, MD) for 1 hr at 55°C. To identify cell types that express miR-544-3p, above tissue sections under ISH were incubated with chicken anti-NeuN antibody (1:200; Aves, Davis, CA) overnight at 4°C followed by donkey anti-chicken IgG Alexa 488 (Jackson ImmunoResearch labs, West Grove, PA) for 1 hr at room temperature. ISH detection was performed using anti-DIG secondary antibody conjugated with radish peroxidase (POD) (Roche Diagnostics Corp., Indianapolis, IN) and Opal 570 (1:500; Perkin Elmer, Waltham, MA). Images were captured using a confocal microscope (Nikon A1+; Nikon Metrology Inc, Brighton, MI). All images were analyzed using NIS elements or ImageJ software in a blinded manner. Three sections of the L4-L5 DRG per animal were analyzed for the quantification of DRG neuron subpopulations.

### DRG neuron culture and transfection

DRG neurons were dissociated and cultured as described previously^32,33^. Briefly, L3-L5 lumbar DRGs from adult mice were harvested and placed in a dish containing complete saline solution (CSS) on ice for cleaning and mincing . CSS contains (in mM): 137 NaCl, 5.3 KCl, 1 MgCl_2_, 3 CaCl_2_, 25 Sorbitol, and 10 HEPES, adjusted to pH 7.2 with NaOH. For 20 min the DRGs were digested with 0.35 U/ml Liberase TM (Roche Diagnostics Corp., Indianapolis, IN) and then for 15 min with Liberase TL (0.25 U/ml; Roche Diagnostics Corp., Indianapolis, IN) and papain (30 U/ml, Worthington Biochemical, Lakewood, NJ) in CSS containing 0.5 mM EDTA at 37°C. The tissue was triturated with a fire-polished Pasteur pipette. After centrifugation, a cell pellet containing 1 x 10^4^ cells was resuspended in 10 μl of buffer R (Thermo Fisher Scientific, Waltham, MA) and DRG neurons were then electroporated with miR-544-3p mimic (50 nM; Thermofisher), or miR-544-3p inhibitor (50 nM; Thermo Fisher Scientific, Waltham, MA), or negative control mimic (50 nM; Thermo Fisher Scientific, Waltham, MA) using a Neon transfection system (Thermo Fisher Scientific, Waltham, MA). Stimulus parameters were set at three pulses (1400 V) of 10 ms duration. The transfected cells were placed onto poly-D-lysine/laminin coated glass coverslips. The complete DRG medium contained equivalent amounts of DMEM and F12 (Gibco, Gaithersburg, MD) with 10% FBS (Hyclone, Logan, UT) and 1% penicillin and streptomycin (Invitrogen, Carlsbad, CA). The cells were then maintained in 5% CO_2_ at 37°C in a humidified incubator.

### Quantitative real-time PCR (qRT-PCR) and bioinformatics analysis

RNA was extracted from L3-L5 DRGs using the RNeasy Lipid Tissue Mini Kit (Qiagen, Germantown, MD). RNA from cultured and transfected DRG cells was purified using the Absolutely RNA Nanoprep Kit (Agilent Tech., Santa Clara, CA). RNA quantity was measured using a NanoDrop 1000 (Thermo Fisher Scientific, Waltham, MA). RNA was then reverse-transcribed to complimentary DNA using the iScript cDNA Synthesis Kit (Bio-Rad, Hercules, CA) according to the manufacturer’s protocol. For relative quantification of mRNA, RT-PCR was performed on a QuantStudio 3 real-time PCR system (Applied Biosciences Corp., Beverly Hills, CA) using PowerUp SYBR Green Master Mix (Applied Biosciences Corp, Beverly Hills, CA). The primer sequences are summarized in Supplemental Table 1. For miR-544-3p, cDNA was reverse-transcribed using a Taqman MicroRNA Reverse Transcription Kit (Appliedbiosystems, Foster City, CA) according to the manufacturer’s protocol. RT-PCR was performed using a MicroRNA Assay Kit (Appliedbiosystems, Foster City, CA) and Taqman Universal Master Mix II (Appliedbiosystems, Foster City, CA). Each sample was assayed in duplicate. The expression levels of the target genes were quantified relative to the level of β-actin (for mRNA) or U6 (for microRNA) gene expression using the 2^-ΔΔCT^ method.

Conserved miRNA target sites on the FcγRI mRNA were predicted using several online programs, including TargetScan (http://www.targetscan.org/), miRDB (http://mirdb.org/miRDB/), and PicTar (http://pictar.mdc-berlin.de/) ^34^. The sequence conservation of miR-544-3p and its binding site in Fcgr1 3’-UTR among rhesus, human, rat, mouse, chimpanzee, and dog was analyzed using online TargetScan software.

### Dual-luciferase reporter-system assays

Wildtype and miR-544-3p binding site-mutated 3’UTR of *Fcgr1* were purchased from Thermo Fisher Scientific (Waltham, MA) and the inserts subcloned into the psiCHECK2 plasmid (Promega, Madison, WI) using GeneArt seamless cloning and assembly enzyme mix (Invitrogen, Carlsbad, CA). HEK293 cells were seeded onto a 24-well culture plate (1 x 10^5^ per well) and cultured overnight. Cells were then co-transfected with recombinant plasmid (300 ng) together with miR-544-3p mimic or negative control mimic (50 nM) using Lipofectamine 3000 (Thermo Fisher Scientific, Waltham, MA) and Lipofectamine RNAimax (Thermo Fisher Scientific, Waltham, MA), respectively. At 48 hrs after transfection, firefly and Renilla luciferase activities were analyzed using Dual-Glo luciferase assays (Promega, Madison, WI). Dual-Glo luciferase reagent was added to each well and firefly luciferase-mediated luminescence was measured using a SpectraMax i3X plate reader (Molecular device, San Jose, CA). Stop & Glo reagent was then added to each well and Renilla luciferase-mediated luminescence was measured. Firefly luminescence was divided by Renilla luminescence, and the resulting luminescence ratio in miR-544-3p-transfected cells was normalized to that in cells transfected with negative control mimic.

### In vivo transfection

In vivo-jetPEI (Polyplus Transfection Inc, New York, NY) was prepared according to the manufacturer’s instructions. A total of 3 µg of miR-544-3p mimic, or miR-544-3p hairpin inhibitor, or negative control (400 µM, Dharmacon) were diluted in a 10 µl solution containing 0.36 µl of in vivo jetPEI and 9.10 µl of 5% glucose. In naïve mice, the mixture was injected intrathecally (i.t.) into mice at L4/5 spinal levels once daily every 4 days for two days. A valid spinal puncture and intrathecal delivery of the mixture was confirmed by a reflexive tail flick after needle entry into the subarachnoid space. At 48 hrs after the last injection, IgG-IC (10 or 100 µg/ml; 10 µl) was injected into one ankle joint of mice. Pain-related behaviors were measured before and 1, 3, 5 hrs after IgG-IC injection. In the CIA model, the mixture of jetPEI in vivo transfection reagent and miR-544-3p mimic or negative control was delivered i.t. into mice once daily on days 42, 45 and 48 after CIA induction. Pain-related behaviors were assessed 72 hrs after each injection.

### Statistical analysis

Data are presented as means ± SEM. A two-tailed Student’s t test was used to test the significance of differences between two groups. Comparisons for multiple groups or multiple time points were carried out using a one-way or two-way ANOVA for random measures or repeated measures followed by Bonferroni’s post hoc test comparisons. P value less than 0.05 was considered significant. Group size and the type of statistical tests used for each comparison are noted in each figure legend. All statistical analyses were performed on GraphPad Prism 7.0 software.

### Data availability

Data that support the findings of this study are available from the corresponding author, upon reasonable request.

## Results

### miR-544-3p is downregulated in mouse DRG in AIA and CIA arthritis models

Our recent study suggested that neuronal FcγRI acts as a crucial driver of arthritis pain^26^. Considering miRNAs as master modulators of gene expression, we utilized several online miRNA target prediction tools (miRDB, TargetScan, and PicTar) to identify a series of potential miRNAs that might target *Fcgr1*. These miRNA candidates included miRNA-6990-3p; miRNA-6909-3p; miRNA-705; miRNA-6993-5p; miRNA-3967; miRNA-6982-3p; miRNA-153-5p; miRNA-7017-3p; miRNA-5626-3p; miRNA-6402; miRNA-7838-3p; miRNA-383-5p, miRNA-544-3p, miRNA-127-3p, miRNA-204-5p, miRNA-378-5p, miRNA-299a-3p, miRNA-299b-3p, miRNA-340-5p, miRNA-133a-3p, miRNA-133b-3p, miRNA-375-3p, miRNA-43-5p, miRNA-133c and miRNA-134-5p. To assess whether these miRNA candidates are related to the development of arthritis pain, we assayed their expression in the DRG in two commonly used animal models of RA using qPCR. Due to high cost of miRNA probes, we designed the primers for these candidates in our screening assay (Supplementary Table 1).

In the AIA model, the expression levels of seven microRNA candidates (miR-127-3p, miR-143-3p, miR-544-3p, miR-378a-5p, miR-133b-3p, miR299a-3p and miR6982) were significantly reduced in the DRG on day 3 after the 2^nd^ challenge (Fig. 1A) whereas the remaining candidates were not significantly changed. In addition, the expression of some miRNA candidates (miR-153-5p, miR-340-5p and miR-3967) was not detectable in the DRG. In the CIA model, the expression levels of miR-127-3p, miR-143-3p, miR-544-3p and miR-7238 were significantly reduced in the DRG on day 56 after CIA induction (Fig. 1B). Among these downregulated FcγRI-targeting microRNA candidates, only miR-127-3p, miR-143-3p and miR-544-3p were downregulated in both AIA and CIA models. Moreover, miR-544-3p was the species with the most robust reduction in the DRG. miR-544-3p was therefore chosen for further anatomical and functional analyses. To further confirm the downregulation of miRNA-544-3p expression in the DRG in the setting of arthritis, we analyzed the time course of its expression during CIA using a Taqman microRNA assay. Downregulation of miRNA-544-3p was observed on days 42 and 56 after CIA but not on day 28 (Fig. 1C). Interestingly, *Fcgr1* mRNA expression was upregulated in the DRG at the same time points, representing a possible inverse correlation with miR-544-3p expression (Fig. 1C). By contrast, this correlation was not observed in spinal cord since the expression of both miR-544-3p and *Fcgr1* mRNA was upregulated in spinal cord on day 56 after CIA induction (Fig. 1D).

**Figure 1.**
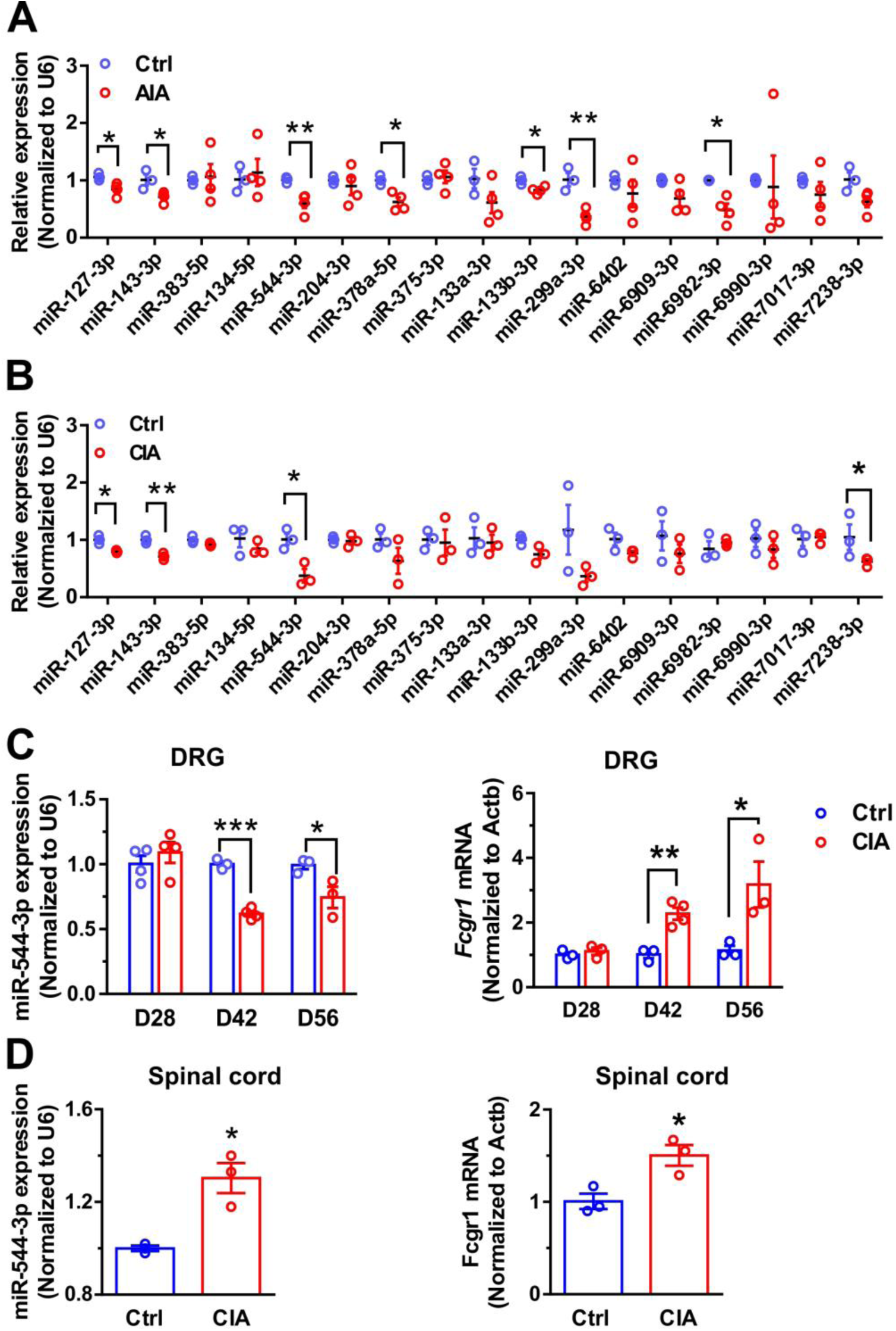
miR-544-3p is downregulated in mouse DRG in AIA and CIA models. (**A, B**) qRT-PCR analysis of FcγRI-targeting miRNA candidate expression (normalized to U6) in L3-L5 DRGs of control (Ctrl) and AIA mice on day 3 after the 2^nd^ challenge (**A**; n = 3–4 mice per group), and control (Ctrl) and CIA mice on day 56 after immunization (**B**; n = 3 mice per group). (**C**) Time course of expression levels of miR-544-3p (left) and *Fcgr1* mRNA (right) in L3-L5 DRGs of control (Ctrl) and CIA mice on days 28, 42 and 56 after immunization (n = 3-4 mice per group). (**D**) qRT-PCR analysis of miR-544-3p (left) and *Fcgr1* mRNA (right) in spinal cord of control (Ctrl) and CIA mice on day 56 after immunization. *p < 0.05, **p < 0.01, ***p < 0.001 versus Ctrl; unpaired Student’s t test without a correction for multiple t-tests.

Next, we defined the expression profile of miR-544-3p in the DRG. In situ hybridization (ISH) analysis revealed miR-544-3p expression in 55% of DRG neurons in naïve mice across a range of soma sizes (Fig. 2A-C). miR-544-3p signal colocalized with immunostaining for the neuronal specific nuclear protein NeuN, suggesting neuronal expression of miR-544-3p (Fig. 2A). The specificity of miR-544-3p detection was validated by a loss of ISH signal in DRG sections stained with a scramble-miR probe (Fig. 2B). On day 56 after CIA induction, a smaller percentage of DRG neurons expressed miR-544-3p compared to that in control mice (Fig. 2D, E). These changes were statistically significant in small and large diameter DRG neurons but exhibited only a nonsignificant trend in medium diameter neurons. (Fig. 2E).

**Figure 2.**
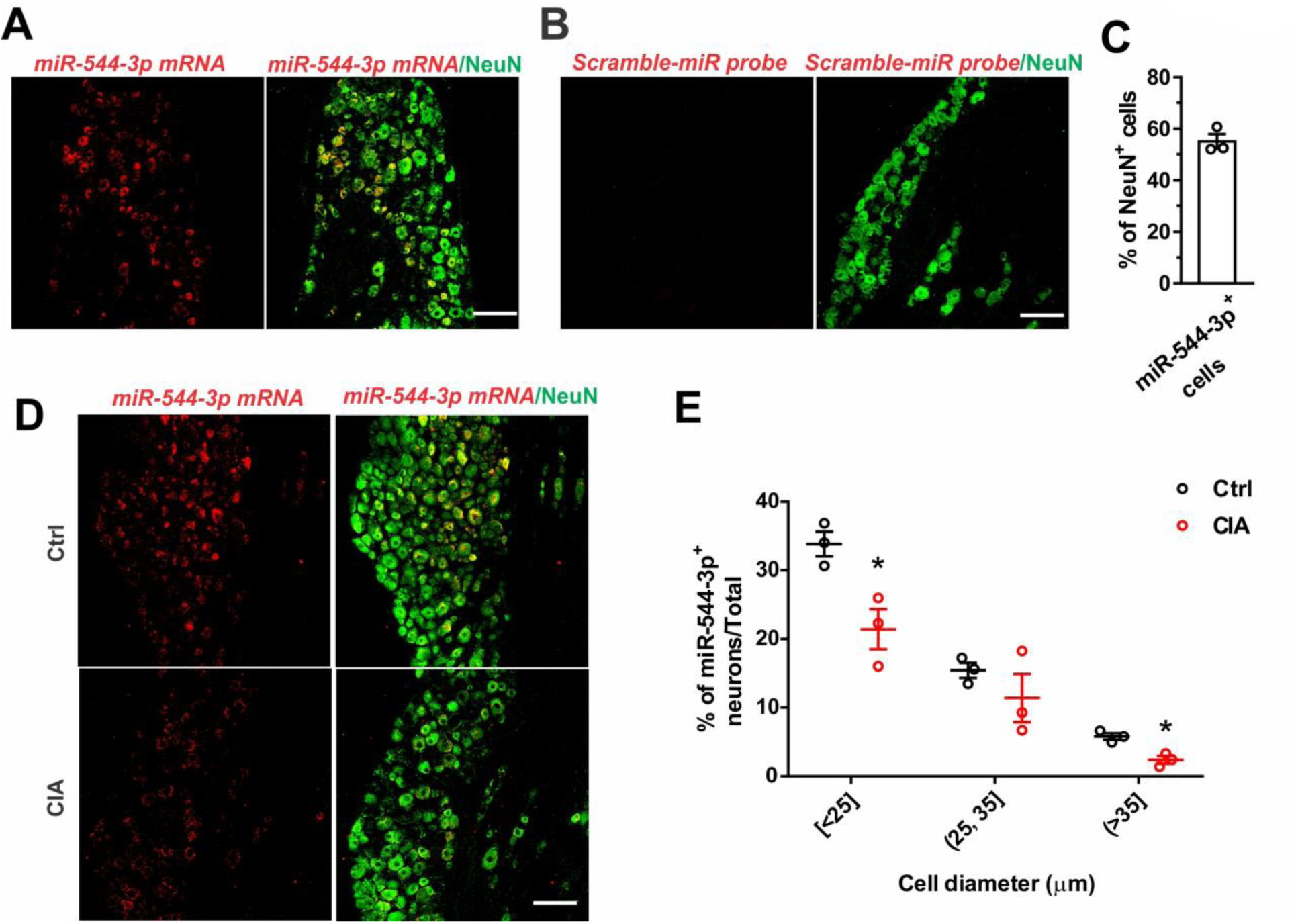
CIA causes a downregulation of miR-544-3p expression in DRG neurons. (**A**) ISH images showing miR-544-3p expression in a subset of DRG neurons of naïve mice (n =3 mice). miR-544-3p signal (red) was colocalized with the pan-neuronal marker NeuN (green). (**B**) ISH images of DRG sections stained with a scramble-miR probe. Scale bar: 100 µm. (**C**) Quantification of miR-544-3p expression in DRG neurons of naïve mice (n = 3 mice). (**D**) ISH images of miR-544-3p expression in the DRG of control (Ctrl) and CIA mice on day 56 after immunization. (**E**) Size frequency distribution of miR-544-3p expression in DRG neurons of control (Ctrl) and CIA mice on day 56 after immunization (n = 3 mice per group). Scale bar: 100 µm. *p < 0.05 versus Ctrl; unpaired Student’s t test.

### miR-544-3p mimic alleviates arthritis pain in the mouse CIA model

To evaluate a potential role of miR-544-3p in the maintenance of arthritis pain, we delivered miR-544-3p mimic or a non-targeting miRNA negative control by i.t. injections into wild-type mice on days 42, 45 and 48 after CIA induction. Mechanical and thermal hyperalgesia were tested 72 hrs after each injection (Fig. 3A). miR-544-3p mimic significantly increased mechanical threshold in the ankle of mice on day 51, compared to the negative control (Fig. 3B). Similarly, mice treated with miR-544-3p mimic exhibited less secondary mechanical and thermal hyperalgesia in the hind paw than those subjected to the negative control on days 45, 48 and 51 (Fig. 3C, D). However, no significant changes in arthritis score were observed between the two groups (Fig. 3E). To explore the potentially direct relationship between miR-544-3p and FcγRI in the context of arthritis pain, we harvested L3-L5 DRGs after behavioral testing on day 51 and found a significant decrease of *Fcgr1* mRNA expression in the DRG but not in spinal cord after miR-544-3p mimic injection compared to negative control injection (Fig. 3F). Of note, miR-544-3p was reported to alleviate neuropathic pain via targeting signal transducer and activator of transcription 3 (STAT3) in spinal cord.^35^ Therefore, we asked whether STAT3 could be another potential target of miR-544-3p in the DRG in the context of CIA. Yet, we did not detect any obvious changes in *stat3* mRNA expression level in the DRG of mice with CIA compared to control mice (Supplementary Fig. 1). To explore whether FcγRI reciprocally regulates the expression of miR-544-3p, we compared the expression level of miR-544-3p in the DRG between *Fcgr1*^+/+^ and *Fcgr1*^−/−^ mice. No significant differences in miR-544-3p expression in the DRG were observed between genotypes (Supplementary Fig. 2). Taken together, these results suggest that miR-544-3p downregulation contributes to the maintenance of arthritis pain, at least in part, through the regulation of FcγRI expression in the DRG. However, we cannot exclude the possibility that miR-544-3p regulates the expression of other genes^35–37^.

**Figure 3.**
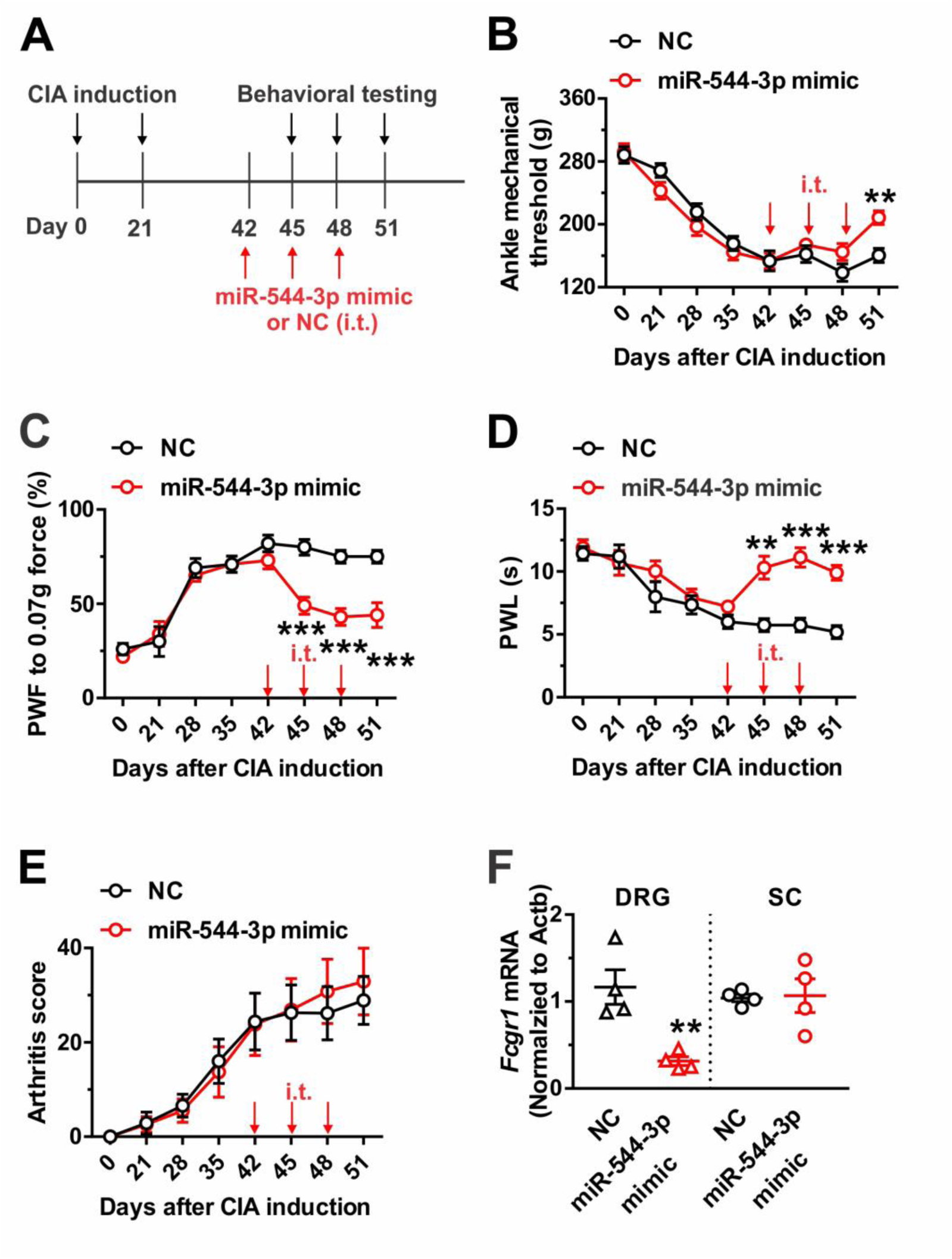
miR-544-3p mimic alleviates arthritis pain by suppressing *Fcgr1* expression in the CIA. (**A**) Intrathecal (i.t.) administration protocol of miR-544-3p mimic (3 µg) or a non-targeting negative control (NC). Pain-like behaviors were evaluated at 72 hrs after each injection. (**B-E)** Time course of mechanical threshold in the ankle (**B**), paw withdrawal frequency (PWF) in response to 0.07g force in the hind paw (**C**), paw withdrawal latency (PWL) to radiant heat in the hind paw (**D**), and arthritis score (**E**) in NC- and miR-544-3p mimic-treated mice with CIA (n = 10 mice per group). **p < 0.01, ***p < 0.001 versus NC; two-way repeated measures ANOVA followed by Bonferroni correction. (**F**) qRT-PCR of *Fcgr1* mRNA expression in L3-L5 DRGs and spinal cord (SC) on day 51 after CIA induction. **p < 0.01; unpaired Student’s t test.

### FcγRI mediates arthritis pain in the CIA model

Although our recent study showed that FcγRI mediated chronic pain in several inflammatory arthritis models^26^, little is known about the potential role of FcγRI in arthritis pain in the CIA model. Assuming that FcγRI is a putative downstream target of miR-544-3p, we hypothesized that *Fcgr1* deletion would produce an analgesic effect in the CIA model similar to that of the miR-544-3p mimic. To test this possibility, we compared pain related behaviors between wildtype (*Fcgr1*^+/+^) and global *Fcgr1* knockout (*Fcgr1^−/−^)* mice following CIA. As expected, we found that *Fcgr1^−/−^* mice displayed less primary mechanical hyperalgesia in the ankle, less secondary mechanical allodynia and thermal hyperalgesia in the hind paw over the course of CIA, compared to wildtype controls (Fig. 4A-C). In addition, the arthritis score in *Fcgr1^−/−^* mice was markedly attenuated throughout the disease stages compared to wildtype controls (Fig. 4D). No obvious gender differences in analgesic effects of *Fcgr1* deletion were observed over the course of CIA (Supplementary Fig. 3).

**Figure 4.**
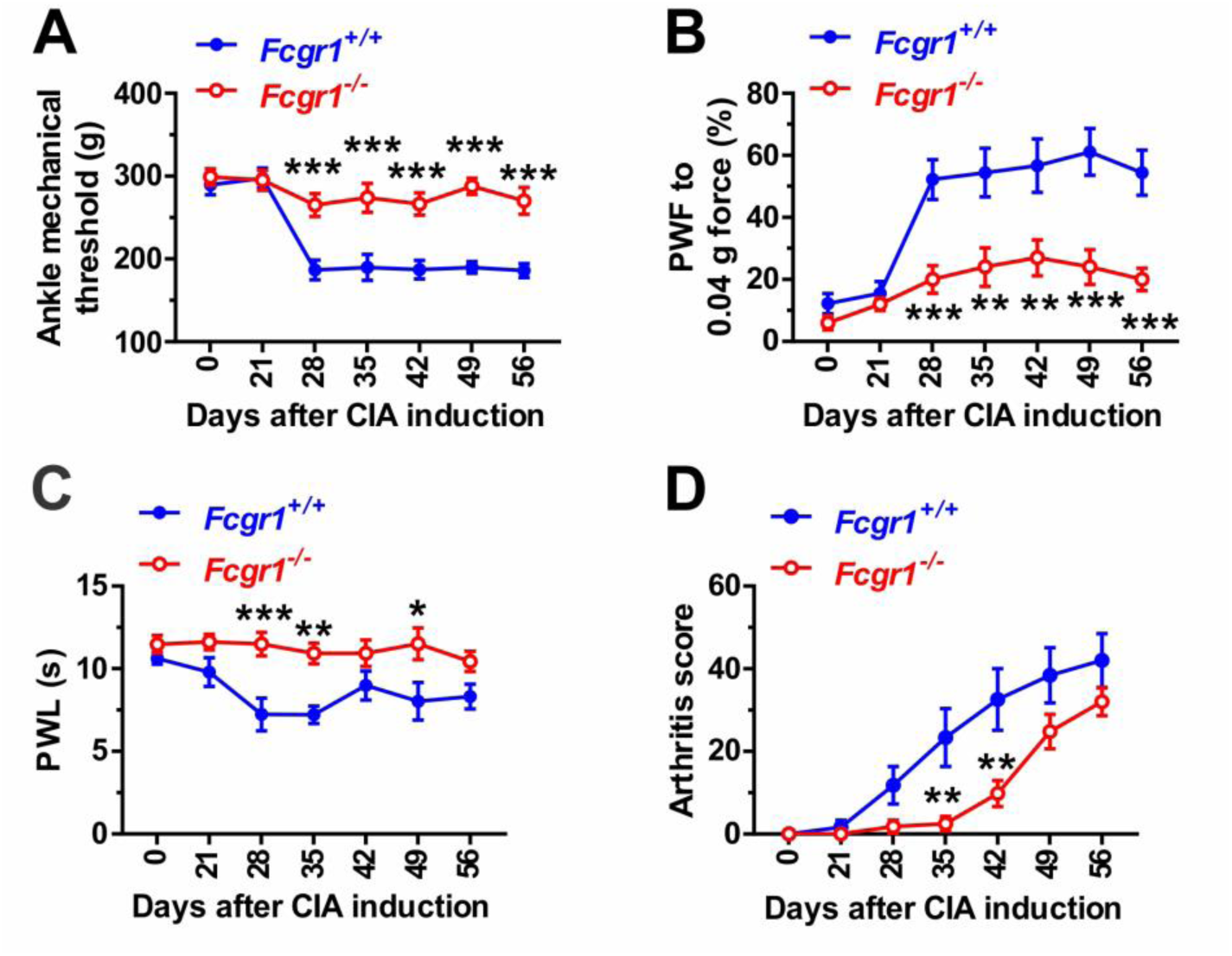
Global genetic deletion of *Fcgr1* attenuates arthritis pain in the CIA model. (**A-D**) Time course of mechanical threshold in the ankle (**A**), paw withdrawal frequency (PWF) in response to 0.04 g force in the hind paw (**B**), paw withdrawal latency (PWL) to radiant heat in the hind paw (**C**), and arthritis score (**D**) in wildtype (*Fcgr1*^+/+^; n = 10 mice) and global *Fcgr1*^-/-^ (n = 11 mice) mice with CIA. * p < 0.05, **p < 0.01; ***p < 0.001 versus *Fcgr1*^+/+^; two-way repeated measures ANOVA followed by Bonferroni correction.

### FcγRI is a direct target of miR-544-3p

The sequence alignments of miR-544-3p are well conserved among mammals, especially within the seed region (Fig. 5A), suggesting that the regulation of FcγRI by miR-544-3p is functionally important. To further assess whether miR-544-3p directly targets FcγRI, we transfected dissociated DRG neurons with miR-544-3p mimic or a negative control. qRT-PCR assay confirmed successful overexpression of miR-544-3p in DRG neurons (Fig. 5B). miR-544-3p mimic significantly reduced *Fcgr1*mRNA expression in DRG neurons compared to the negative control (Fig. 5C). By contrast, transfection of miR-544-3p inhibitor induced an upregulation of the expression of *Fcgr1*mRNA in DRG neurons (Fig. 5D), suggesting that FcγR is a potential target of miR-544-3p and that its expression is under tonic suppression by this miRNA. Given that the 3’-UTR of genes is a site commonly bound by miRNAs^38^, we used an online bioinformatics algorithm (TargetScan; http://www.targetscan.org/) to predict and locate potential binding sites for miRNA-544-3p in *Fcgr1* 3’-UTR (Fig. 5E).

**Figure 5.**
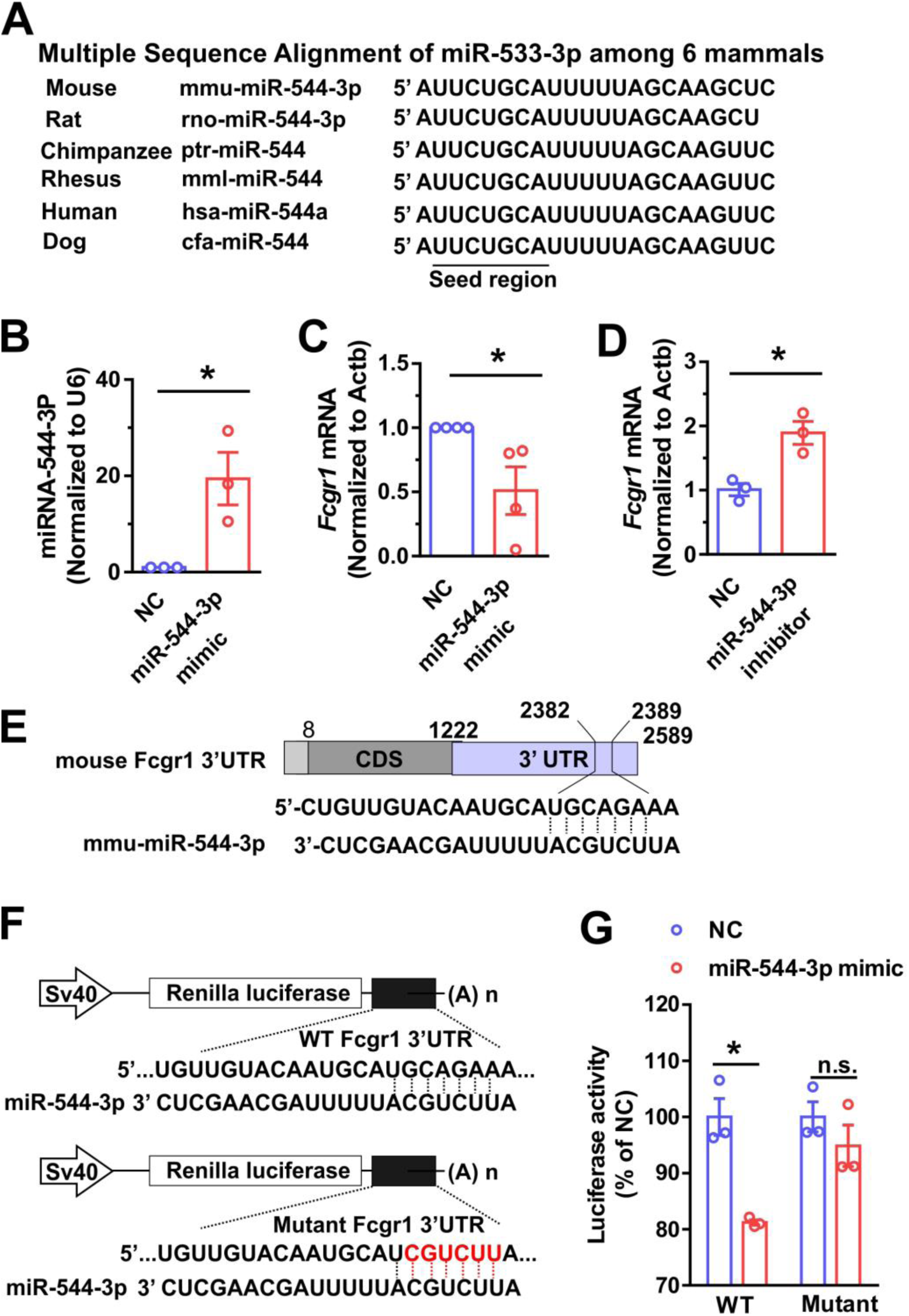
miR-544-3p directly targets the *Fcgr1* 3’UTR. (**A**) Sequence alignment of miR-544-3p in different mammals. Data was acquired at miRbase database (http://www.mirbase.org/). **(B-D)** qRT-PCR analysis of miR-544-3p (**B**) and *Fcgr1* expression (**C, D**) in dissociated DRG neurons electroporated with miR-544-3p mimic (50 nM) or miR-544-3p inhibitor (50 nM) or a negative control (50 nM) at 48 hrs after transfection; n = 3 samples; *p< 0.05 versus NC; unpaired Student’s t test. (**E**) Predicted binding position of miR-544-3p on Fcgr1 3’UTR. (**F**) Schematic representation of the psiCHECK2 constructs containing the full length of wildtype (WT) or mutant Fcgr1 3’UTR. The mutant sequence is indicated by red letters. (**G**) The luciferase activity of WT *Fcgr1* -3’UTR-psicheck2 (WT) or mutant -*Fcgr1-* 3’UTR-psiCheck2 (Mutant) in HEK293 cells cotransfected with miR-544-3p mimic or a negative control (NC). n = 3 samples *p< 0.05 versus NC; unpaired Student’s t test.

To further confirm whether miR-544-3p directly targets the 3’-UTR of *Fcgr1*, we constructed luciferase reporter vectors (psiCHECK-2) containing either the full-length wildtype *Fcgr1* 3’-UTR (WT-Fcgr1-3’UTR-psiCHECK-2) or a miR-544-3p binding site mutant *Fcgr1* 3’-UTR (Mutant-Fcgr1-3’UTR-psiCHECK-2) and analyzed the effect of miR-544-3p on luciferase activity using a dual-luciferase assay (Fig. 5F). We cotransfected WT-Fcgr1-3’UTR-psiCHECK-2 or Mutant -Fcgr1-3’UTR-psiCHECK-2 together with miR-544-3p mimic or a negative control into HEK293 cells. Transfection of miR-544-3p mimic resulted in a significant reduction of luciferase activity in HEK293 cells transfected with WT-Fcgr1-3’UTR-psiCHECK-2 but not in those transfected with Mutant-Fcgr1-3’UTR-psiCHECK-2 (Fig. 5G), indicating that miR-544-3p can downregulate *Fcgr1* mRNA expression by directly binding to the predicted site of *Fcgr1* 3’UTR.

### miR-544-3p overexpression attenuates acute joint pain hypersensitivity induced by IgG-IC via FcγRI

Our recent study showed that IgG-IC was sufficient to evoke acute joint pain hypersensitivity through FcγRI in naïve mice^26^. Given that miR-544-3p is able to target and downregulate FcγRI, we postulated that miR-544-3p would modulate IgG-IC evoked nocifensive responses by targeting FcγRI. To test this possibility, we analyzed whether upregulation of miR-544-3p reduces pronociceptive effects of IgG-IC. A total of 3 µg miR-544-3p mimic or a non-targeting negative control was mixed with jetPEI in vivo transfection reagent (10 µl) and injected i.t into wildtype mice once daily every 4 days for two days. At 48 hrs after the last injection, IgG-IC (100 µg/ml; 10 µl) was injected into one ankle joint (Fig. 6A). Pain-related behaviors were measured before and 1, 3, and 5 hrs after IgG-IC injection. miR-544-3p mimic had no significant effects on basal mechanical sensitivity in the hind ankle or hind paw compared to miRNA controls (Fig. 6B, C). However, miR-544-3p mimic significantly alleviated primary mechanical hyperalgesia in the ankle and secondary mechanical hyperalgesia in the hind paw elicited by i. a. injection of IgG-IC (Fig. 6B, C). We harvested the DRG and spinal cord immediately after behavioral testing. qRT-PCR analysis revealed an obvious decrease in *Fcgr1* mRNA expression and a robust increase in miR-544-3p expression in both DRG and spinal cord after i.t. injection of miR-544-3p mimic compared to the negative control (Fig. 6D, E). ISH analysis further confirmed that *Fcgr1* mRNA signal in DRG neurons was suppressed by miR-544-3p mimic (Fig. 6F, G). These results suggest that exogenous miR-544-3p attenuates IgG-IC evoked articular hypernociception, at least in part, by regulating FcγRI expression in DRG neurons.

**Figure 6.**
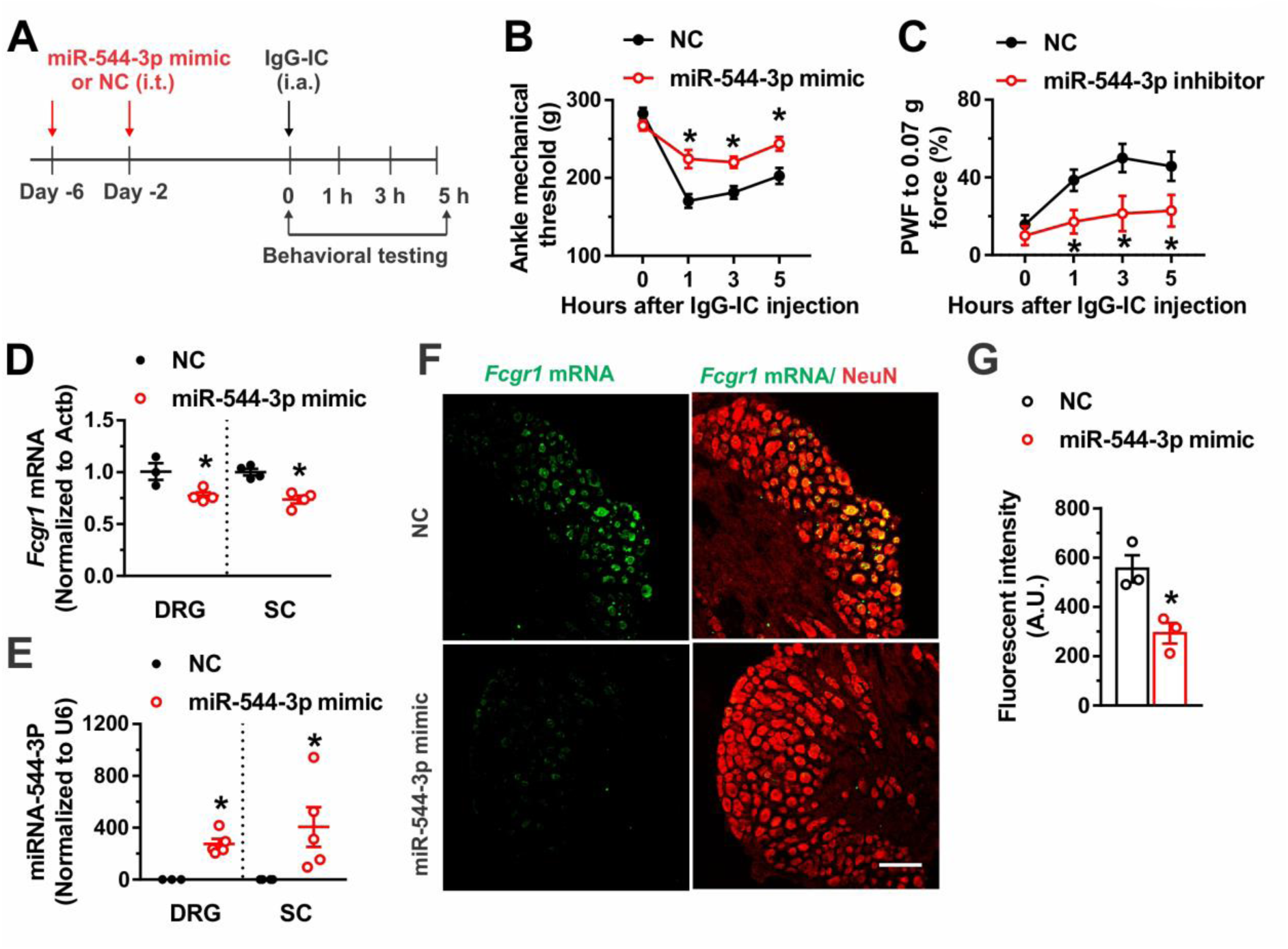
miR-544-3p mimic alleviates IgG-IC induced acute joint nocifensive behaviors via the downregulation of FcγRI expression in naïve mice. (**A**) Intrathecal (i.t.) administration protocol of miR-544-3p mimic (3 µg) or a non-targeting negative control (NC). At 48 hrs after the last miRNA injection, mice were injected intraarticularly (i.a.) with IgG-IC (100 µg/ml; 10 µl) and pain-like behaviors were evaluated over 1-5 hrs. (**B, C)** Mice treated with miR-544-3p mimic exhibited higher mechanical threshold in the ankle (**B**) and lower paw withdrawal frequency in response to 0.07 g force (**C**) applied to the hind paw following i.a injection of IgG-IC compared to those treated with NC. n = 7 mice/group. Arrow indicates the time of injection of miR-544-3p or NC. *p < 0.05 versus NC; two-way repeated measures ANOVA followed by Bonferroni correction. (**D, E)** qRT-PCR analysis of the expression of *Fcgr1* mRNA normalized to that of *Actb* and miR-544-3p normalized to U6 in the DRG (**D**) and spinal cord (SC; **E**) of NC-(n = 3 mice) and miR-544-3p mimic treated mice (n = 5 mice). * p < 0.05 versus NC; unpaired Student’s t-test. (**F**) Representative L4 DRG in situ hybridization image for *Fcgr1* and immunostaining for NeuN from NC- and miR-544-3p mimic-treated mice. Scale bar: 100 µm. (**G**) Quantification shows significant reductions of *Fcgr1* mRNA expression in DRG neurons of miR-544-3p mimic-treated mice compared with NC-treated animals (n = 3 mice per group). * p < 0.05 versus NC; unpaired Student’s t-test.

### miR-544-3p inhibitor potentiates acute joint pain hypersensitivity induced by IgG-IC

Next, we assessed whether inhibition of miR-544-3p potentiates IgG-IC induced nocifensive behaviors via upregulation of FcγRI. We injected i.t. a 10 µl mixture of miR-544-3p inhibitor (3 µg) or a negative control (3 µg) with jetPEI in vivo transfection agent into naïve mice once daily every four days for two days. At 48 hrs after the last injection, IgG-IC (10 µg/ml; 10 µl) was injected to one ankle of mice (Fig. 7A). No obvious differences in basal mechanical sensitivity at the hind ankle or hind paw skin were observed between negative control- and miR-544-3p inhibitor-treated mice (Fig. 7B, C). However, primary mechanical hypersensitivity in the ankle and secondary mechanical hypersensitivity in the hind paw evoked by i.a. injection of IgG-IC were significantly enhanced in mice treated with miR-544-3p inhibitor compared to the negative control (Fig. 7B, C). qRT-PCR analysis revealed that miR-544-3p inhibitor caused a small but significant increase in *Fcgr1* mRNA expression in the DRG (Fig. 7D). Yet, no significant differences in *Fcgr1* mRNA expression in spinal cord were observed between the two groups (Fig. 7D).

**Figure 7.**
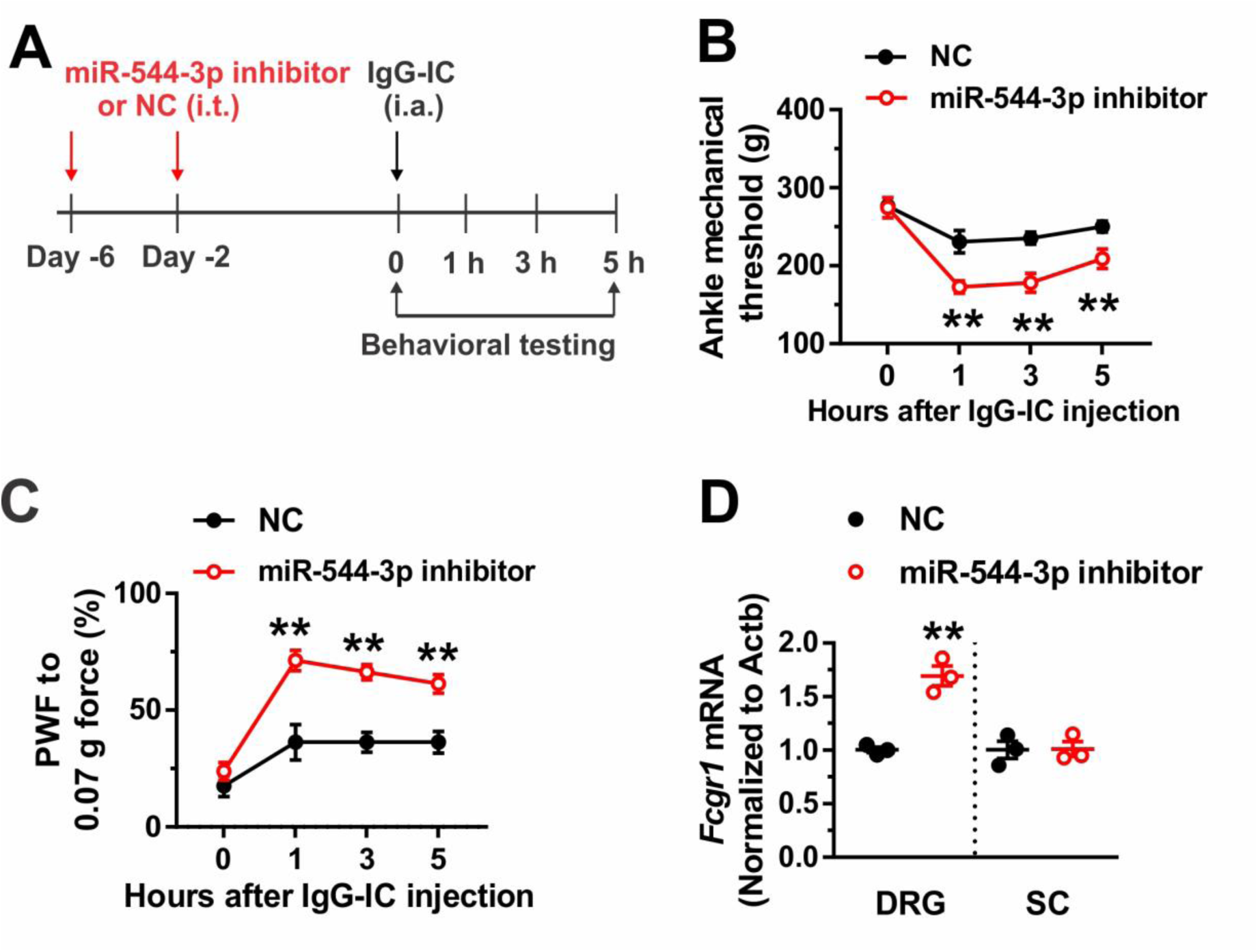
miR-544-3p inhibitor enhances acute joint pain hypersensitivity induced by IgG-IC in naïve mice. (**A**) Experimental schematic showing intrathecal (i.t.) administration protocol of miR-544-3p mimic (3 µg) or a negative control (NC) as well as the time for behavioral testing after intraarticular (i.a.) injection of IgG-IC (10 μg/ml; 10 μl). (**B, C**) Time course of mechanical threshold of the ankle joint and paw withdrawal frequency (PWF) in response to 0.07 force applied to the hind paw in NC- and miR-544-3p- treated mice after i.a injection of IgG-IC. n = 8 mice per group. *p< 0.05, **p < 0.01 versus NC; two-way repeated measures ANOVA followed by Bonferroni correction. (**D**) qRT-PCR analysis of the expression of *Fcgr1* mRNA normalized to that of actb in the DRG (**D**) and spinal cord (SC; **E**) of NC- and miR-544-3p mimic treated mice (n = 3 mice per group). *p < 0.05 versus NC; unpaired Student’s t test.

## Discussion

As master regulators of pain gene expression, miRNAs have emerged as critical pain modulators ^14,15^. miR-544-3p is widely expressed in a variety of cell types and has been implicated in multiple physiological and pathological processes, such as cancer, stroke and cardiovascular disease ^37,39,40^. Here, we have provided novel evidence that miR-544-3p is also expressed in primary sensory neurons and acts as a key functional miRNA for the maintenance of arthritis pain in mouse RA models. Our qRT-PCR and ISH analyses revealed that the expression of miR-544-3p was downregulated in joint-innervating DRG in the context of arthritis. Rescuing miRNA-544-3p downregulation remarkably attenuated arthritis pain in the CIA model. The function of miR-544 in the central nervous system has been widely studied ^40–42^. The expression of miR-544a was downregulated after spinal cord injury in mice and overexpression of miR-544a alleviated neuroinflammation in the spinal cord and restored motor function^42^. In addition, miR-544-3p in the spinal cord was implicated in the pathogenesis of neuropathic pain^35^. In the rat CCI model, miR-544-3p expression was downregulated in the dorsal horn and overexpression of miR-544-3p in spinal cord alleviated neuropathic pain and reduced proinflammatory cytokines expression in CCI rats ^35^. In the present study, we extended previous findings by showing that miR-544-3p may possess analgesic effects on arthritis pain at the periphery. Given that miR-544-3p mimic was administered through intrathecal delivery and that the upregulation of miR-544-3p expression were observed in both DRG and the spinal cord of naïve mice after i.t administration, we cannot completely rule out a central action of miR-544-3p mimic. Unlike neuropathic pain conditions, we found that miR-544-3p expression was downregulated in the DRG while upregulated in spinal cord in the setting of arthritis. Thus, it is possible that the observed analgesic effect of miR-554-3p mimic on arthritis pain is mainly mediated by a peripheral mechanism.

FcγRI is typically expressed in immune cells and plays a key role in the pathogenesis of RA^43^. In addition to immune cells, we and other groups have identified that FcγRI is also expressed in primary sensory neurons^25,26,44,45^. Moreover, our recent study showed that neuronally expressed FcγRI mediated hyperactivity of joint sensory neurons and arthrits pain in animal models of RA through a mechanism independent of inflammation^26^. Although proinflammatory cytokines may contribute to the upregulated FcγRI expression in DRG neurons during arthritis, it is also possible that FcγRI expression is controlled by epigenetic modifications^46^. In the present study, serval complementary lines of evidence suggest that FcγRI is a target of miR-544-3p in the DRG. First, the expression of miR-544-3p was downregulated while the mRNA expression of *Fcgr1* was upregulated in the DRG in AIA and CIA models. The inverse correlation between the expression of FcγRI and miR-544-3p is consistent with miR-544-3p targeting FcγRI. By contrast, we did not observe such inverse correlation in spinal cord since both molecules were upregulated in the CIA model. Therefore, it is possible that reversal of miR-544-3p targeting and regulation of FcγRI occurs mainly in the DRG in the setting of arthritis. Second, in the CIA model, overexpression of miR-544-3p reduced *Fcgr1* mRNA expression in the DRG and attenuated arthritis pain. Accordingly, genetic deletion of FcγRI produced antihyperalgesic effects similar to those of miR-544-3p mimic. Considering that FcγRI is a key immune receptor expressed in a variety of immune cells, adeno-associated virus (AAV) delivery itself may cause the upregulation of FcγRI expression by mobilizing immune cells. Due to this limitation, our study could not evaluate whether AAV vector-mediated FcγRI overexpression could abolish antihyperalgesic effects of miR-544-3p mimic. Third, in dissociated DRG neurons, *Fcgr1* mRNA expression was downregulated by miR-544-3p mimic and upregulated by miR-544-3p inhibitor. Fourth, bioinformatics algorithms predicted that the 3’UTR of Fcgr1 possesses a match to the seed sequence of miR-544-3p. Using the dual-luciferase assay, we found that miR-544-3p mimic reduced luciferase reporter activity in HEK293 cells transfected with a construct containing the full length of 3’UTR sequence of *Fcgr1* but not one containing a mutant *Fcgr1*-3’UTR. These findings further confirmed that miR-544-3p directly targets the 3’UTR of *Fcgr1* to downregulate *Fcgr1*mRNA expression. Lastly, in naïve mice, miR-544-3p mimic downregulated *Fcgr1* mRNA expression in the DRG and alleviated IgG-IC-induced joint pain hypersensitivity. By contrast, miR-544-3p inhibitor upregulated *Fcgr1* mRNA expression in the DRG and potentiated IgG-IC-induced nocifensive behaviors. Given that FcγRI is a receptor for IgG-IC and mediates pronociceptive effects IgG-IC^26^, we suggest that miR-544-3p contributes to IgG-IC-induced acute joint pain hypersensitivity, at least in part, through the regulation of *Fcgr1* mRNA expression. Unlike under arthritis conditions, the downregulation of *Fcgr1* mRNA expression by miR-544-3p mimic was observed in both DRG and spinal cord in naive mice. We therefore cannot completely exclude a central action of miR-544-3p in naïve mice.

Each miRNA is able to target hundreds of genes, and each mRNA can also be targeted by multiple miRNAs. In this study, we found that the expression of several other FcγRI targeting miRNA candidates besides miR-544-3p were also downregulated in the DRG under arthritis conditions, including miR-127-3p and miR-143-3p. Since miR-544-3p exhibited the most robust downregulation in the DRG, the present study mainly focused on this candidate. A recent study also showed the downregulation of miR-143-3p expression in the DRG in the CIA model^22^. Moreover, miR-143-3p negatively regulated the expression of certain pain associated genes in DRG neurons, such as *ptgs2*, *mrgpre* and *tnf* ^22^. In addition, the miR-127-3p/FcγRI axis has been well documented in the setting of lung inflammation^46^. Further studies are needed to define the contribution of miR-143-3p and miR-127-3p to RA pain. Although the present study identified that FcγRI is a direct target of miR-544-3p in the DRG during arthritis, we cannot rule out the possibility that other miR-544-3p-targeted genes are also involved in miR-544-3p-induced analgesia. miR-544-3p was reported to attenuate neuropathic pain by downregulating the expression of STAT3 in spinal cord ^35^. However, we did not observe any significant changes in *stat3* mRNA expression in the DRG in the context of arthritis, suggesting that STAT3 may not be a direct of target of miR-554-3p in the DRG under arthritis conditions. In cancer tissues, miR-544-3p functions as a tumor suppressor by directly targeting a variety of genes, such as ras homolog family member A (RHOA) and Toll-like receptor 2 (TLR2) ^36,37,47^. Further investigation is required to explore whether any of these potential targets mediate analgesic effects of miR-544-3p in the setting of arthritis.

Mature miRNAs act as posttranscriptional regulators of gene expression, by base-pairing with mRNA molecules in the 3’-UTR or controlling the translation process of the target mRNA ^48^. In the present study, *Fcgr1* mRNA expression was downregulated by miR-544-3p in DRG neurons, suggesting that miR-544-3p posttranscriptionally controls FcγRI expression, possibly via mRNA degradation. Since all commercially available anti-FcγRI antibodies we tested lacked adequate specificity for reliable western blotting and immunohistochemical staining, it was infeasible to test whether miR-544-3p can suppress FcγRI translation. We cannot completely rule out the possibility that miR-544-3p downregulates FcγRI expression via translational repression. In addition to their conventional role as gene regulators, miRNAs can also directly interact with proteins in a non-canonical manner^49^. miR-let-7b was shown to directly activate TLR7/TRPA1 in nociceptive neurons to trigger acute pain hypersensitivity ^50^. Another recent study identified that miR-21 functions as a ligand for TLR-7 and contributes to the development of osteoarthritis pain ^51^. Future studies are required to assess whether a similar non-canonical mechanism is involved in the analgesic effect of miR-544-3p.

The cellular and molecular mechanisms underlying miR-544-3p downregulation remain elusive. Bidirectional modulatory networks between miRNAs and their target genes represent a possible mechanism for the regulation of miRNAs. The expression of TGF-β and miR-30c-5p was reported to be bidirectionally regulated in spinal cord in the context of neuropathic pain^52^. In the present study, we showed that miR-544-3p downregulated *Fcgr1* mRNA expression. However, genetic deletion of *Fcgr1* had no effects on miR-544-3p expression in the DRG. It is therefore unlikely that FcγRI directly regulates miR-544-3p expression in the DRG. The downregulation of miR-544-3p expression has been observed in diverse cell types and conditions^35,53^. miR-544 expression was downregulated in spinal cord in a rat model of CCI. Moreover, miR-544 appears to be a direct target of X inactivate-specific transcript (XIST), a long non-coding RNA. XIST contributes to neuropathic pain development in rats through downregulating miR-544^35^. In cervical cancer tissues, Krueppel-like factor 4 (KLF4), a transcription factor, was involved in transcriptional regulation of miR-544-3p through interaction with the miR-544 promoter^53^. Given that both XIST and KLF4 are expressed in the DRG^54,55^, further investigation is required to investigate whether miR-544-3p expression can be regulated by these two molecules under arthritic conditions.

In conclusion, our results provide novel evidence that miR-544-3p contributes to the development of RA pain. Moreover, this analgesic action appears to occur at least in part through the regulation of FcγRI expression in primary sensory neurons. We suggest that miR-544-3p may represent a new candidate therapeutic target for the treatment of chronic pain associated with RA or other autoimmune diseases.

## Acknowledgements

We thank Dr. Sjef Verbeek (Leiden University Medical Center, Leiden, Netherlands) for providing the donor breeders of global Fcgr1 knockout mice. We also thank Ian Reucroft and John Robinson for mouse colony maintenance.

## Supplementary material

Supplementary material is available at Brain online

**Abbreviations**
AAV: adeno-associated virus
AIA: Antigen-induced arthritis
CCI: chronic construction injury
CFA: Complete Freund’s adjuvant
CIA: collagen II-induced arthritis
CSS: complete saline solution
DIG: digoxigenin
DRG: dorsal root ganglion
i.a.: intraarticular
IFA: incomplete Freund’s Adjuvant
IgG-IC: IgG-immune complex
IHC: immunohistochemistry
ISH: in situ hybridization
i.t.: intrathecal
KLF4: Krueppel-like factor 4
mBSA: methylated bovine serum albumin
miRNAs: microRNAs
Mrgpre: MAS related G protein coupled receptor family member E
PFA: paraformaldehyde
Ptgs2: prostaglandin-endoperoxide synthase 2
PWF: paw withdrawal frequency
PWL: paw withdrawal latency
qRT-PCR: Quantitative real-time PCR
RA: rheumatoid arthritis
RHOA: ras homolog family member A
STAT3: signal transducer and activator of transcription 3
s.c.: subcutaneous
TLR2: Toll-like receptor 2
TNF: tumor necrosis factor
UTR: untranslated regions
XIST: X inactivate-specific transcript

