## Supplemental file for "miR-544-3p mediates arthritis pain through regulation of FcγRI"

**Supplemental Table 1. List of primer sequences for quantitative real-time PCR**

| **Genes** | **Sequences 5'-3'** |
| --- | --- |
| mmu-U6-RT | AAAATATGGAACGCTTCACGAATTTG |
| mmu-U6-Foward | CTCGCTTCGGCAGCACATATACT |
| mmu-U6-Reveres | ACGCTTCACGAATTTGCGTGTC |
| Common miR-Reverse | GTGCAGGGTCCGAGGT |
| mmu-miR-127-3p -RT | GTCGTATCCAGTGCAGGGTCCGAGGTATTCGCACTGGATACGACAGCCAA |
| mmu-miR-127-3p -Forward | CGTCGGATCCGTCTGAGC |
| mmu-miR-143-3p -RT | GTCGTATCCAGTGCAGGGTCCGAGGTATTCGCACTGGATACGACGAGCTA |
| mmu-miR-143-3p -Forward | CGCGTGAGATGAAGCACTG |
| mmu-miR-204-5p-RT | GTCGTATCCAGTGCAGGGTCCGAGGTATTCGCACTGGATACGACAGGCAT |
| mmu-miR-204-5p -Forward | CGCGTTCCCTTTGTCATCCT |
| mmu-miR-378a-5p-RT | GTCGTATCCAGTGCAGGGTCCGAGGTATTCGCACTGGATACGACACACAG |
| mmu-miR-378a-5p -Forward | CGCTCCTGACTCCAGGTC |
| mmu-miR-383-5p-RT | GTCGTATCCAGTGCAGGGTCCGAGGTATTCGCACTGGATACGACAGCCAC |
| mmu-miR-383-5p -Forward | CGCGAGATCAGAAGGTGACT |
| mmu-miR-544-3p -RT | GTCGTATCCAGTGCAGGGTCCGAGGTATTCGCACTGGATACGACGAGCTT |
| mmu-miR-544-3p -Forward | CGCGATTCTGCATTTTTAGC |
| mmu-miR-375-3p -RT | GTCGTATCCAGTGCAGGGTCCGAGGTATTCGCACTGGATACGACTCACGC |
| mmu-miR-375-3p -Forward | CGTTTGTTCGTTCGGCTC |
| mmu-miR-299a-3p-RT | GTCGTATCCAGTGCAGGGTCCGAGGTATTCGCACTGGATACGACAAGCGG |
| mmu-miR-299a-3p-Forward | CGCGTATGTGGGACGGTAAA |
| mmu-miR-299b-3p-RT | GTCGTATCCAGTGCAGGGTCCGAGGTATTCGCACTGGATACGACGGTTTA |
| mmu-miR-299b-3p-Forward | CGCGTATGTGGGACGG |
| mmu-miR-340-5p-RT | GTCGTATCCAGTGCAGGGTCCGAGGTATTCGCACTGGATACGACAATCAG |
| mmu-miR-340-5p -Forward | CGCGCGTTATAAAGCAATGAGA |
| mmu-miR-133a-3p-RT | GTCGTATCCAGTGCAGGGTCCGAGGTATTCGCACTGGATACGACCAGCTG |
| mmu-miR-133a-3p-Forward | CGTTTGGTCCCCTTCAAC |
| mmu-miR-133b-3p-RT | GTCGTATCCAGTGCAGGGTCCGAGGTATTCGCACTGGATACGACTAGCTG |
| mmu-miR-133b-3p-Forward | CGTTTGGTCCCCTTCAAC |
| mmu-miR-133c-RT | GTCGTATCCAGTGCAGGGTCCGAGGTATTCGCACTGGATACGACCTGACT |
| mmu-miR-133c-Forward | CGTTTGGTCCCCTTCAAGG |
| mmu-miR-134-5p-RT | GTCGTATCCAGTGCAGGGTCCGAGGTATTCGCACTGGATACGACCCCCTC |
| mmu-miR-134-5p-Forward | CGTGTGACTGGTTGACCA |
| mmu-miR-6990-3p -RT | GTCGTATCCAGTGCAGGGTCCGAGGTATTCGCACTGGATACGACCTGCCA |
| mmu-miR-6990-3p -Forward | CGAGCCCTGCCTCTTCC |
| mmu-miR-6909-3p -RT | GTCGTATCCAGTGCAGGGTCCGAGGTATTCGCACTGGATACGACCTGTGG |
| mmu-miR-6909-3p -Forward | TGCCTTCCCCGGCCTC |
| mmu-miR-6993-5p -RT | GTCGTATCCAGTGCAGGGTCCGAGGTATTCGCACTGGATACGACGACAGA |
| mmu-miR-6993-5p -Forward | CGAGTGGGAGAAACGGGTG |
| mmu-miR-3967-RT | GTCGTATCCAGTGCAGGGTCCGAGGTATTCGCACTGGATACGACCAACAT |
| mmu-miR-3967 -Forward | CGCGAGCTTGTCTGACTG |
| mmu-miR-6982-3p -RT | GTCGTATCCAGTGCAGGGTCCGAGGTATTCGCACTGGATACGACCTGGAA |
| mmu-miR-6982-3p -Forward | TGGCCCCTCTGCCCC |
| mmu-miR-153-5p -RT | GTCGTATCCAGTGCAGGGTCCGAGGTATTCGCACTGGATACGACAGCTGC |
| mmu-miR-153-5p -Forward | CGCGGTCATTTTTGTGACGTT |
| mmu-miR-7017-3p-RT | GTCGTATCCAGTGCAGGGTCCGAGGTATTCGCACTGGATACGACCTGGAG |
| mmu-miR-7017-3p-Forward | CGACCCTGCTCCTCTCC |
| mmu-miR-5626-3p-RT | GTCGTATCCAGTGCAGGGTCCGAGGTATTCGCACTGGATACGACGTGTCA |
| mmu-miR-5626-3p-Forward | CGCAGCAGTTGAGTGATG |
| mmu-miR-6402-RT | GTCGTATCCAGTGCAGGGTCCGAGGTATTCGCACTGGATACGACGCGGGT |
| mmu-miR-6402-Forward | CGGAGCAGTTTTCCCAGGA |
| mmu-miR-7238-3p-RT | GTCGTATCCAGTGCAGGGTCCGAGGTATTCGCACTGGATACGACGAACAC |
| mmu-miR-7238-3p-Forward | CGCGCGCGCTGTCCTTT |
| Fcgr1-Forward | TGCTGGATTCTACTGGTGTGA |
| Fcgr1-Reverse | AAACCAGACAGGAGCTGATGA |
| Stat3-Forward | AGCTGGACACACGCTACCT |
| Stat3-Reverse | AGGAATCGGCTATATTGCTGGT |
| Actb-Forward | CTGAATGGCCCAGGTCTGA |
| Actb-Reverse | CCCTGGCTGCCTCAACAC |

RT: Reverse Transcription

**Supplemental Figure legends**

**Supplemental Figure 1. STAT3 expression is not changed in the DRG in the CIA.** qRT-PCR analysis of *Stat3* mRNA in L3-L5 DRGs of control (Ctrl) and CIA mice on days 42 and 56 after immunization. n = 3-4 mice per group. P > 0.05 versus Ctrl, unpaired Student’s t-test.

**Supplemental Figure 2. Genetic deletion *Fcgr1* does not alter miR-544-3p expression in the DRG.** qRT-PCR analysis of miR-544-3p expression in the DRG of *Fcgr1+/+* and *Fcgr1–/–* mice. n = 3 mice per group. P > 0.05 versus Ctrl, unpaired Student’s t-test.

**Supplemental Figure 3. No apparent sex differences were observed in analgesic effects of Fcgr1 deletion in the CIA** (**A-F**) Time course of mechanical threshold in the ankle (**A, D**), paw withdrawal frequency (PWF) in response to 0.04 g force in the hind paw (**B, E**), paw withdrawal latency (PWL) to radiant heat in the hind paw (**C, F**) in male (n = 5 mice per group) and female (n = 5-6 mice per group) wildtype and global *Fcgr1-/-* mice with CIA. * p < 0.05, **p < 0.01; versus *Fcgr1+/+*; two-way repeated measures ANOVA followed by Bonferroni correction.

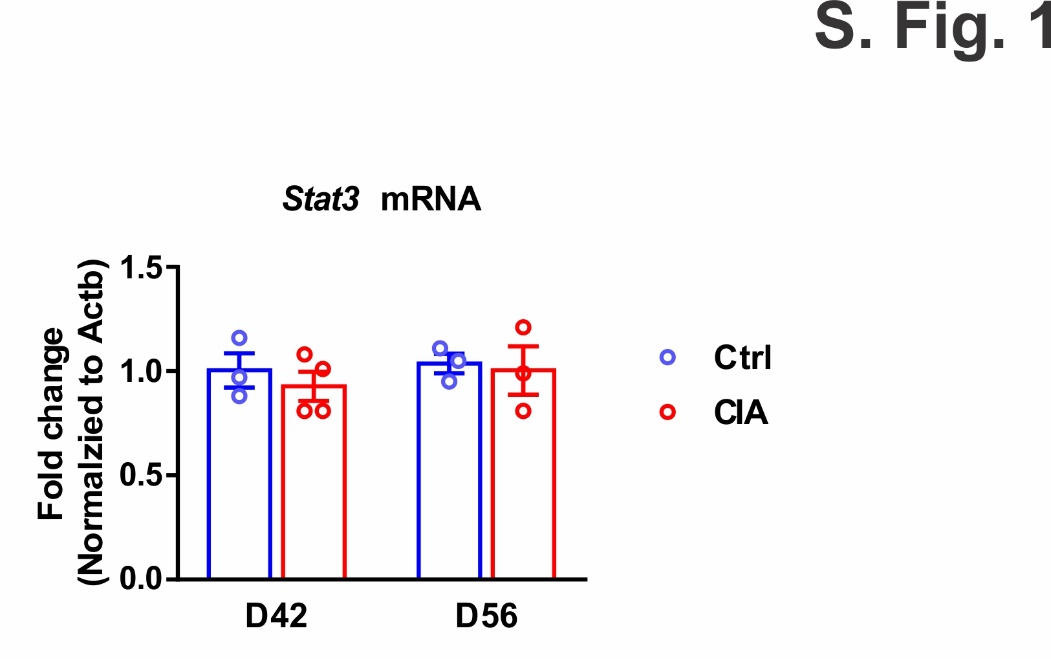

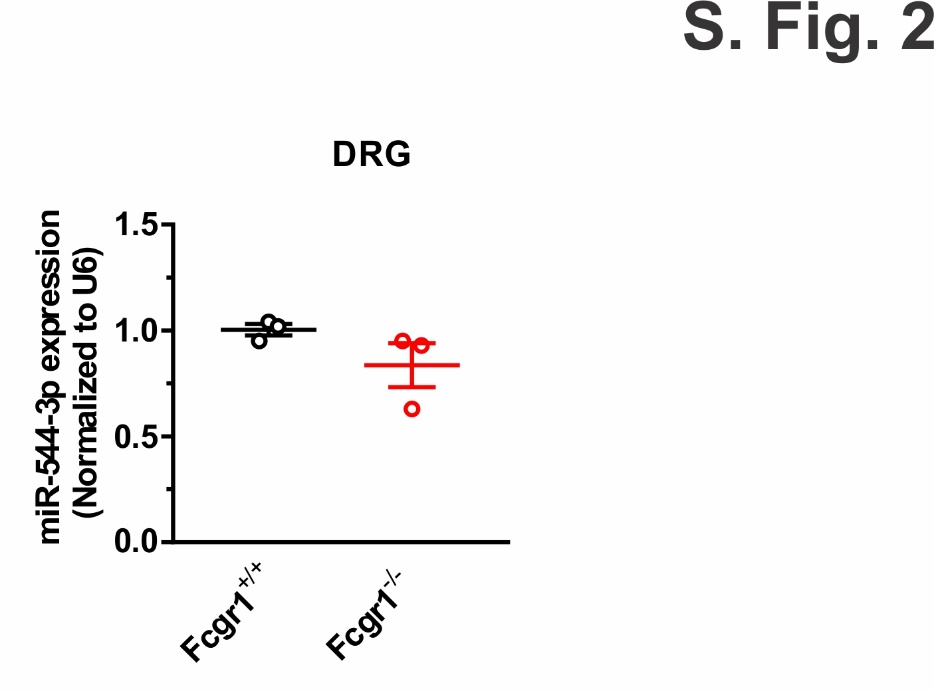

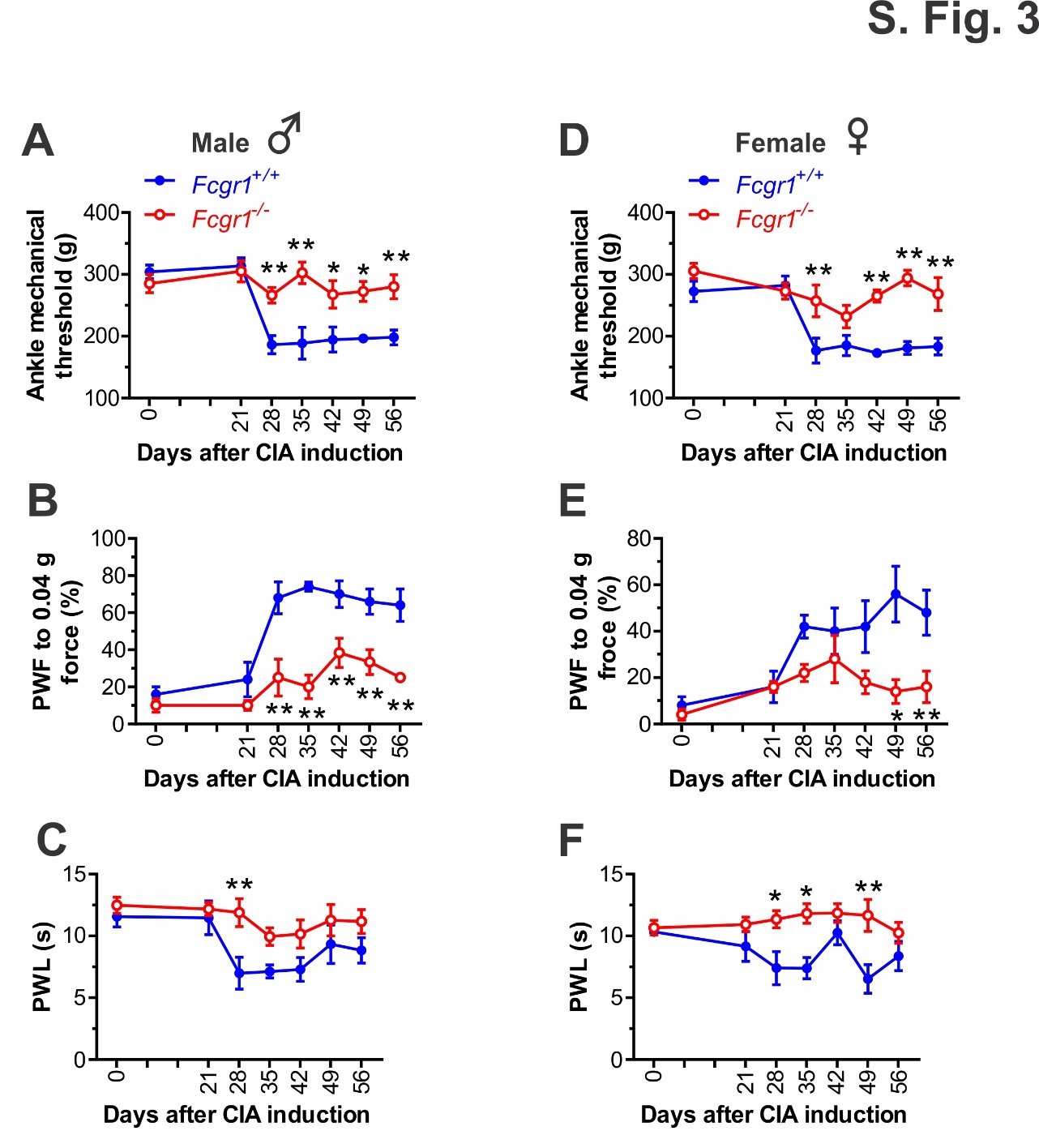
